# Building memories on prior knowledge: behavioral and fMRI evidence of impairment in early Alzheimer’s Disease

**DOI:** 10.1101/2020.06.17.154104

**Authors:** Pierre-Yves Jonin, Quentin Duché, Elise Bannier, Isabelle Corouge, Jean-Christophe Ferré, Serge Belliard, Christian Barillot, Emmanuel J. Barbeau

**Affiliations:** Centre de recherche Cerveau et Cognition, (CerCo), CNRS UMR 5549, Université de Toulouse Paul Sabatier, Toulouse, France; Univ Rennes, Inria, CNRS, INSERM, IRISA, Empenn ERL U-1228, F-35000 Rennes, France; CHU Rennes, Neurology Department, F-35033 Rennes, France; CHU Rennes, Radiology Department, F-35033 Rennes, France

**Keywords:** Alzheimer’s Disease, Prior Knowledge, Source Memory, Recognition Memory, Perirhinal Cortex, fMRI

## Abstract

Impaired memory is a hallmark of prodromal Alzheimer’s Disease (AD). Prior knowledge associated with the memoranda has proved to have a powerful effect on memory in healthy subjects. Yet, barely nothing is known about its effect in early AD. We used functional MRI to ask whether prior knowledge enhanced memory encoding in early AD and whether the nature of prior knowledge mattered. Early AD patients and healthy controls underwent a task-based fMRI experiment, being scanned while learning face-scene associations. Famous faces carried Pre-Experimental Knowledge (PEK) while unknown faces repeatedly familiarized prior to learning carried Experimental Knowledge (EK). As expected, PEK increased subsequent memory in healthy elderly. However, patients did not benefit from PEK. Partly non-overlapping brain networks supported PEK vs. EK encoding in healthy controls. Patients displayed impaired activation in a right subhippocampal region where activity predicted successful associative memory formation of PEK stimuli. These findings call for a thorough consideration of how prior knowledge impacts learning and suggest a possible underestimation of the extent of associative memory impairment in early AD.

**Highlights:**

- Learning is impaired in prodromal AD, but we currently ignore whether prior knowledge available at encoding promotes learning in AD as it does in healthy controls.
- Patients with AD failed to benefit from pre-experimental prior knowledge (famous faces) by comparison with experimental knowledge (unknown but familiarized faces).
- fMRI responses at study revealed distinct networks underlying associative encoding for both pre-experimental and experimental knowledge.
- A subsequent memory effect found in control subjects for associations carrying pre-experimental knowledge in the right subhippocampal structures, including the perirhinal cortex, was absent in patients.
- Pre-experimental knowledge-based associative encoding relies on brain regions specifically targeted by early tau pathology.
- Using unfamiliar materials to probe memory in early AD might underestimate learning impairment.

## 1. Introduction

The idea that prior memories may alter new learning can be tracked back to Hermann Ebbinghaus. While he attempted to rigorously investigate human learning, his use of meaningless syllables was purposeful, as he did not want new learning to be corrupted by things he already knew. Far from Plato’s metaphor of the wax tablet (Roediger, 1980), new learning rarely occurs in a vacuum: instead and typically, new learning is also processed in relation to existing knowledge. Years of research have now shown that prior knowledge about the memoranda has a very powerful effect on memory (e.g. Fernández and Morris, 2018; Umanath and March, 2014). For example, it is easier to remember meeting a neighbor in a given setting than to remember an unknown face, even presented repeatedly. Surprisingly, very little is known about the influence of prior knowledge in Alzheimer’s disease. In other words, it is unknown whether prior knowledge is as effective at promoting memory in AD patients as in healthy subjects and whether the neural correlates of this effect are altered in this disease.

In experimental settings, the ability of participants to remember stimuli carrying “pre-experimental prior knowledge” (PEK), such as famous faces, is often contrasted with their ability to remember either completely unknown items, or stimuli carrying “experimental prior knowledge” (EK), i.e. resulting from repeated exposures to initially unknown stimuli, such as unknown faces (Poppenk et al., 2010a). Evidence from behavioral studies has robustly supported the role of PEK in enhancing associative learning in healthy adults (Bird et al., 2011; Ellis et al., 1979; Greve et al., 2007; Klatzky and Forrest, 1984; Long and Prat, 2002; Reder et al., 2013), including the elderly (Badham et al., 2015; Badham and Maylor, 2015; McGillivray and Castel, 2010).

Task-based functional MRI studies in young adults have highlighted the role of the medial temporal lobes and temporal poles, together with the ventromedial and inferior / middle prefrontal cortex in PEK-based learning when compared to either novel stimuli or recently learned stimuli (Leveroni et al., 2000; Liu et al., 2016; Van Kesteren et al., 2012). EK-based learning in contrast has been associated with activity in the parietal regions and the posterior hippocampus (Dennis et al., 2015; Poppenk et al., 2010a, 2010b; Poppenk and Norman, 2012).

In this study, we aimed at assessing whether PEK benefits memory encoding of patients at the early stage of AD and whether the neural correlates of PEK- vs. EK-based memory encoding are similarly impaired by the disease. For that purpose, we asked patients with early AD (patients with Mild Cognitive Impairment due to Alzheimer’s Disease, “AD-MCI”) and matched healthy controls to encode face-scene associations using an event-related fMRI task design. Prior knowledge about the stimuli was manipulated using either famous faces (PEK condition) or unknown faces the subjects were familiarized with just before the learning phase (EK condition). To identify the brain areas involved in the encoding task, we used the neural adaptation effect, i.e. modulation of the Blood-Oxygen-Level-Dependent (BOLD) fMRI response due to repetition of the stimuli (Grill-Spector et al., 2006).

## 2. Material and methods

### 2.1. Participants

The study was approved by a national ethics committee and is registered in the Clinical Trials database (EPMR-MA Study 2014-A01123-44). Twenty-two patients fulfilling the NIA-AA criteria for Mild Cognitive Impairment due to Alzheimer’s disease (AD-MCI) (Albert et al., 2011) together with 25 healthy controls were screened to participate. Patients were recruited as part of the “Centre Mémoire de Ressources et de Recherches de Haute-Bretagne” at Rennes University Hospital, a Memory Clinic with over 20 years of clinical expertise in the field, and diagnosed by a senior neurologist (SB). Inclusion criteria for AD-MCI patients were: i) evidence of a concern regarding a change in cognition; ii) impaired episodic memory confirmed through neuropsychological assessment; iii) fully preserved independence in functional abilities; iv) evidence for hippocampal atrophy; v) evidence for amyloidopathy either through cerebrospinal fluid (CSF) measures of lower Ab42 levels or via positron-emission tomography (PET) evidence of Ab deposition. Further inclusion criteria were i) age 60-75 years; ii) >7 years of education; iii) French native speaking; iv) right-handedness. Exclusion criteria were i) any history of alcoholism, drug abuse, head trauma or psychiatric condition; ii) 7-items modified Hachinski ischemic score>2 (Hachinski et al., 2012); iii) scores above age- and gender-adjusted available cut-off at the Beck Depression Inventory (BDI-II, Beck et al., 1996) or at the State-Trait Anxiety Inventory (STAI, A&B, Spielberger et al., 1983); iv) dementia (Mckhann et al., 2011).

All subjects underwent two testing sessions. They first underwent an extensive neuropsychological assessment (see Supplementary Materials for details) which allowed to (1) rule out any subtle cognitive impairment among healthy controls; (2) rule out severe impairments in AD-MCI patients that would be incompatible with the second experimental session; (3) avoid the inclusion of atypical Alzheimer’s disease profiles like particular progressive focal degenerative phenotypes among our experimental group (Alladi et al., 2007). The second visit included the imaging and behavioral experimental sessions.

Overall, five AD-MCI were finally excluded from the original sample (two presented with severely impaired cognition preventing them from undergoing the experiments, one scored above the cut-off at the depression inventory, one gave-up during the second visit, and one was discovered to suffer claustrophobia in the scanner). Six healthy controls were also excluded (two due to technical issues during MRI acquisition, one due to cognitive scores below norms, one due to back pain complaint in the scanner, one due to above cut-off score at the depression inventory, and one due to the discovery of a pituitary adenoma), resulting in the final inclusion of 19 healthy controls and 17 AD-MCI.

### 2.2. Design & general procedure

Once the first testing session was completed, participants came back to the lab within one month for the second session. The whole procedure of the second session is illustrated in **Figure 1**. It was divided in four sequential phases: 1) familiarization with the MRI environment and with a series of unknown faces in a mock scanner; 2) study phase (i.e., learning) during real MRI acquisition; 3) recognition memory test outside the scanner and 4) fame judgment for all items involved in the study phase.

**Figure 1.**
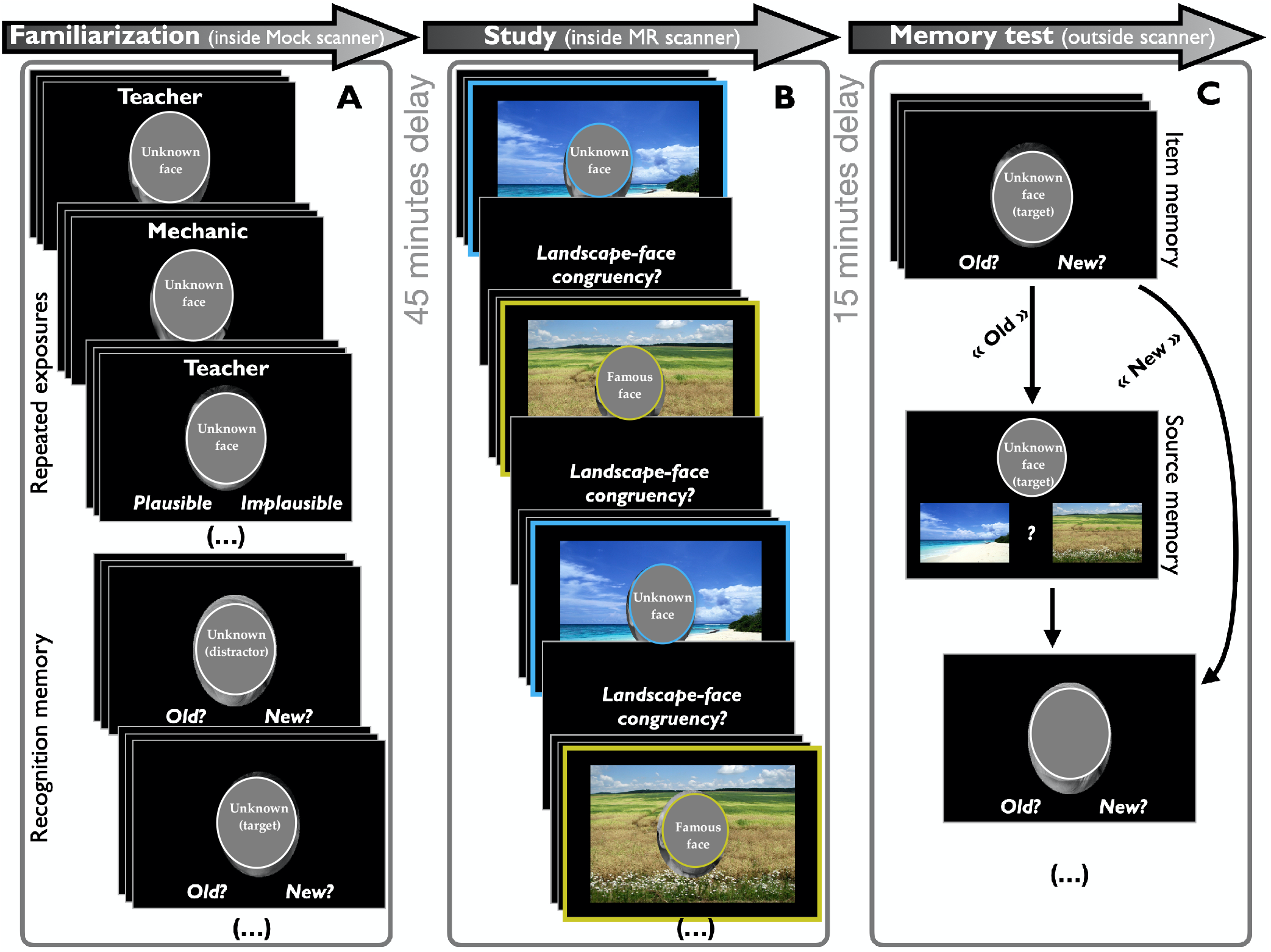
Experimental design. Photographs of faces were obscured due to the editor policy. **(A)** During the **Familiarization phase**, unknown faces were repeatedly presented inside a mock scanner. Participants were instructed to make a congruency judgment for the face-occupation association. Then, an immediate Old/New recognition test was administered. **(B)** Forty-five minutes later, **the study phase** inside the MR scanner involved the explicit encoding of face-scene associations of two types: EK trials (blue outline, here only for illustrative purpose), involved a face from the familiarization phase; PEK trials (yellow outline) involved a famous face. The “Landscape-face congruency?” also displayed photos of the handgrips the participants used to provide their response (not depicted). **(C)** The **Memory test phase** took place outside the scanner after a 15 minutes delay. Participants had to make Old/New judgments for individual faces. For each Hit (i.e. correct “Old”) response, a two-alternate forced choice test asked subjects to recall the correct source (i.e. which scene was associated with the face at study). Finally, after a 5-minutes break, a **Fame judgment** test (not depicted) involved the whole set of famous and unknown faces.

### 2.3. Cognitive tasks

#### Stimuli

During the “study phase” performed in the MRI scanner, unique associations between a scene (landscape picture) and a face were used as stimuli (**Figure 1**). Famous faces were used in the PEK condition, while unknown faces (familiarized through repeated presentations before the encoding phase) were used in the EK condition. Face images were converted to greyscale pictures and cropped to a 250 px-width oval-shape. Extensive pre-testing of fame judgment in 35 healthy elderly (aged 55-79, not included in the present study) resulted in the selection of two sets of n = 132 famous faces and n = 184 unknown faces, that were matched as closely as possible for sex ratio, age, ethnicity, hair colors, emotional expression and paraphernalia. Ninety-eight colored landscape images were selected so that half represented a beach and half a countryside. None included humans, animals nor artifacts. They were all normalized to 720×484 px. Two sets of 48 famous and 48 unknown faces were chosen at random, each being randomly associated with 24 countryside and 24 beach landscapes images, resulting in 96 PEK stimuli (i.e. famous face - landscape associations) and 96 EK stimuli (i.e. unknown face - landscape associations), half being used as targets and half as distractors. An additional set of 26 unknown faces was randomly chosen for use as distractors in the recognition test during the familiarization phase (see below & **Figure 1**). The order of presentation, the set of pictures as well as the face-scene combinations were fully randomized across participants.

#### Familiarization phase (pre-scan)

Participants were installed in a mock MRI scanner designed to familiarize them with the real MRI scanner environment, including space, noise, luminosity, handgrip used as the response device, and computer screen viewed through a rear-facing mirror. They were presented a series of faces randomly associated with an occupation, and were instructed to make a congruency judgment about that association. Forty-eight unknown faces were repeatedly presented 3 times across 6 blocks, with face-occupation combinations remaining constant across repetitions. For each participant, a pseudo-random order of presentation was used so that at least 3 trials separated two identical stimuli. Each stimulus was presented for 2.5 seconds followed by a 1.5 second time response window. No memory instruction was given. Immediately after that familiarization phase, participants were administered a surprise Old/New recognition memory test involving the 48 target faces together with 26 new unstudied, unknown distractor faces. The purpose of this testing session was to assert that each participant had correctly encoded the 48 unknown target faces. Following the recognition test, participants remained in the mock scanner and received the instructions and some practice trials for the real fMRI study phase.

#### Study phase (inside the MRI scanner)

Forty-five minutes after the familiarization phase, participants underwent the study task, inside the MRI scanner. The critical trials required subjects to explicitly learn face-scene associations. Each stimulus was presented for 3.5 seconds, then the participant had 1.5 second to decide whether the background scene was congruent or not with the face. The congruency task was designed to ensure that enough attention was paid to both the face and the scene at encoding, and participants were explicitly instructed that they would be further tested for their memory of the associations. The arrow task (Stark and Squire, 2001) was adapted and used as an active baseline, and jittered white fixation crosses were displayed between trials. The schedule of events was optimized via Optseq2 (Dale, 1999). Each run started and ended with a 4-seconds fixation cross and included 44 events: 16 EK, 16 PEK and 12 Arrow. In total, 48 EK and 48 PEK trials were repeated once so that each study event was presented twice across the 6 runs, without any repetition within a run. Median lag between two identical events was 8 minutes. Lag durations between two occurrences of the same stimulus were similar for PEK and EK events (670 vs. 630 seconds for EK & PEK, respectively, U=5035; *p*=0.268). The order in which runs were presented was counterbalanced across participants.

#### Test phase (outside the scanner)

Fifteen minutes after the study phase, participants were administered a recognition memory test in a quiet room. Target faces were randomly mixed with distractors and participants had to make an Old/New decision by reference to the study phase. This provided a measurement of item memory, i.e. memory for faces only. After Hit responses (i.e. “Old” responses to a target face), the face was presented again together with two scenes, one featuring a beach and the other one a countryside. Participants were instructed to choose the correct source, namely the scene associated with the face at study. Since both scenes were taken from the study phase, absolute familiarity alone could not trigger the correct choice. This provided a behavioral proxy for associative memory accuracy. The order of presentation was fully randomized across participants.

#### Fame judgment phase

Five minutes after the test phase, subjects were shown all the faces from the test phase again and asked to make a “Famous/Unknown” judgment. As a result, any PEK or EK stimulus yielding inaccurate responses at that step was removed from further analyses. This allowed us to contrast truly famous (i.e. items associated with PEK) vs. truly unknown (i.e. items associated with EK due to the familiarization phase only) on an individual basis. The participants provided accurate responses, ranging from 77 to 100%. Importantly, the control and patient groups did not differ regarding their fame judgment performance, with similar accuracy across all items (U=132; *p*=0.357), also when considering more specifically the PEK faces (U=122; *p*=0.215).

### 2.4. Behavioral data analysis

Item recognition memory performance indices (i.e. memory for faces only, Hits and False Alarm rates) were computed within the signal detection theory framework. Following Verde et al., (2006), *A_z_* was computed to estimate sensitivity, namely, how well participants discriminated between targets and distractors. Accordingly, we computed a non-parametric metric of bias *B”* (Grier, 1971). These indices were preferred to the parametric *d’* and *C* indices of sensitivity and bias, respectively, for their superior robustness to the underlying assumptions regarding responses distributions (Stanislaw and Todorov, 1999; Verde et al., 2006), and corresponding formulae were implemented using a dedicated Excel workbook (Gaetano, 2017; Signal detection theory calculator 1.2 [Excel workbook downloaded from https://www.researchgate.net/profile/Justin_Gaetano2]).

Associative learning performance was estimated through source memory accuracy. Source memory refers to the ability to correctly recall the context that was associated with the target item at study. Here, we measured source memory as the conditional probability of a Source Hit (i.e. giving a correct source response) given an Item Hit (i.e. giving a correct “Old” response to a target face), which is a classical behavioral proxy for associative memory accuracy (Cooper et al., 2017). Repeated-measures ANOVAs were run to explore whether the kind of prior knowledge (EK vs. PEK) altered sensitivity (item memory), bias, associative memory (item + context memory), both within and between groups. Parametric statistical testing was used when the assumptions of normality and variance equality were met. Otherwise, non-parametric methods were used. Analyses were performed using the JASP software (https://jasp-stats.org, JASP Team (2018). JASP (Version 0.9) [Computer software]).

### 2.5. Structural and functional imaging

#### 2.5.1. Image acquisition

Participants were scanned using a 3T Verio (Magnetom Siemens, Siemens Healthcare, Erlangen, Germany) system equipped with a 32-channels phased-array coil running VB17. High-resolution (1mm isotropic) MPRAGE T1-weighted images were collected (TR/TI/TE = 1900/900/2.26ms, FOV 256×256, 176 slabs, parallel imaging (GRAPPA) factor 2) for anatomical visualization and spatial normalization. Blood-oxygen level dependent (BOLD) functional images were collected using a T2*-weighted single-shot spin-echo EPI sequence with the following parameters: repetition time = 2,000 ms, echo time = 30 ms, 3×3×3.6 mm^3^ voxel size, 192×192 mm^2^ field-of-view, 64×64 matrix, slice thickness = 3.6 mm, 36 slices, parallel imaging (GRAPPA) factor 2, echo-spacing 0.51 ms, bandwith 2368Hz/Px, spacing between slices = 0.6 mm. A total of 840 volumes divided in 6 sessions (runs) of 140 volumes were acquired for each participant. Each session lasted 4 minutes and 40 seconds. The task was run using the E-Prime 2.0 software (Psychology Software Tools, Pittsburgh, PA). A rear-facing mirror allowed participants to see the stimuli on an LCD screen placed at the back of the scanner, and they gave their responses using a two-button response handgrip (NordicNeurolab, Bergen, Norway).

#### 2.5.2. fMRI data preprocessing

Image preprocessing was performed using SPM12 (7219, https://www.fil.ion.ucl.ac.uk/spm/). For each participant, a subset of BOLD images was randomly selected for visual checking. Functional images were then corrected for slice acquisition temporal delay and spatially realigned to the across-run mean image to correct for subject motion. Then, they were coregistered to the T1-weighted anatomical image and normalized to the Montreal Neurological Institute (MNI) stereotactic space at a 2×2×2 mm³ resolution before being spatially smoothed using an 8×8×8 mm^3^ full width at half maximum Gaussian kernel.

#### 2.5.3. fMRI data analysis

For each participant, a general linear model (GLM) was estimated voxel-wise. The experimental design for the individual statistical analysis was modeled with thirteen regressors: a 3×2×2 factorial design plus a regressor for the active baseline condition. The regressors of interest referred to prior knowledge status (EK, PEK), to repetition (the order of events presentation, i.e. first “p1”, second “p2”), and to subsequent memory (Source Hit (SH), Source Miss (SM, i.e. Item Hit followed by incorrect source response) and Miss (M, i.e. Item Miss, that is, incorrect “New” response to a target face)). Events of interest (face-scene associations) were modeled with 3.5 seconds boxcar functions and a 3s boxcar function was used to model the active baseline task, but null events (i.e. jittered fixation) were not modeled (Pernet, 2014; Stark and Squire, 2001). The regressors of interest and the active baseline were convolved with the canonical hemodynamic response function. Head motion (6 parameters estimated during the realignment pre-processing step) and magnetic field drift were added as confounding factors. At the group level, contrast images from the subject-level analyses were used to perform a one-sample t-test and evaluate the contrasts of interest group-wise. Two sample t-tests were also performed to probe differences between groups. A individual voxel threshold of p<0.005 uncorrected was used with a cluster extent threshold of 57 contiguous voxels to correct for multiple comparisons (FWE) at *p*<0.05 at the cluster level. This cluster size extent was computed using Monte Carlo simulations (N=10,000 iterations) (Slotnick, 2017; Slotnick et al., 2003). For a recent example of a similar thresholding approach, see (Thakral et al., 2017).

This factorial design setup allowed to estimate four main contrasts used to perform our analyses of interest detailed below. The “Encoding” contrast corresponded to all our regressors of interest minus the active baseline. The “Repetition” contrast corresponded to the subtraction between the first and second presentation of all our regressors of interest. The “Prior Knowledge” contrast corresponded to the subtraction between PEK and EK events of interest. Finally, we estimated the “Prior Knowledge x Repetition” interaction contrast: {PEKp1 – PEKp2 – EKp1 + EKp2}, where PEK and EK included SH, SM and M regressors for each type of prior knowledge. Our analysis workflow thereafter involved the following three steps.

***First***, we investigated whether repetition effects allowed us to identify the brain networks involved in explicit encoding of our stimuli. It is well acknowledged that repetition of stimuli can result in decreased (“Repetition Suppression”) or increased (“Repetition Enhancement”) of the BOLD signal, in brain areas consistent with the ongoing processing, an observation also referred to as “Neural adaptation” (Grill-Spector et al., 2006). Neural adaptation has been successfully used to map functional brain networks, notably memory encoding (Rand-Giovannetti et al., 2006; Reggev et al., 2016) and especially face encoding (Henson, 2016; Henson et al., 2002). We therefore performed a conjunction analysis between the Encoding and Repetition contrasts, to further confirm that our repetition design would highlight common activations within the bilateral visual ventral pathways. ***Second***, as we expected repetition effects to highlight a visual associative encoding network, we computed the [Prior Knowledge x Repetition] interaction contrast to test the hypothesis that prior knowledge modulates brain activity specifically related to face-scene associative encoding. Clusters identified through this contrast were further explored with repeated-measures ANOVAs on the extracted beta weights. ***Third***, beta-weights corresponding to the memory regressors were extracted within the above-defined clusters to further look for subsequent associative memory effects. For that purpose, repeated-measures ANOVAs were computed with the following 4 regressors of interest: EK.SHp1; EK.SMp1; PEK.SHp1; PEK.SMp1. Here, we focused on whether SH and SM differed for EK and PEK. Importantly, only beta weights associated with the first occurrence of the above-mentioned regressors were taken into account for the subsequent memory analysis, thus avoiding confusion between memory and repetition effects, and keeping the interaction and subsequent memory analyses orthogonal. One can refer to (Reggev et al., 2016) for a recent similar approach coupling repetition and subsequent memory effects.

## 3. Results

### 3.1. Behavioral results

#### 3.1.1. Neuropsychological and AD biomarkers findings

AD-MCI patients and healthy controls were matched for age, gender, education, and premorbid verbal IQ, but the patients’ mean MMSE score was, as expected, significantly lower (**Table 1**). Detailed neuropsychological and psychological background of the participants is provided in **Supplementary Material 1** (**Table S.1.)**. AD-MCI essentially presented impairments on recall and recognition memory tests compared to healthy controls. Measurements of hippocampal volumes in both groups confirmed atrophy in the patient group (Manjón and Coupé, 2016). Biomarker investigations confirmed amyloidopathy in patients (Abeta42 dosage in CSF, n = 7, or abnormal amyloid retention using 18F-AV-45-PET, n = 7; data not available for 3 patients) (**Table 1**). Our AD-MCI group therefore fulfills the research diagnostic criteria for AD as the etiology of their cognitive impairments (Albert et al., 2011).

**Table 1.** Demographic, clinical and AD biomarkers characteristics of the participants. (*TIV= Total Intracranial Volume)

|  | Healthy Controls | AD-MCI | p values |
| --- | --- | --- | --- |
| N | 19 | 17 | - |
| Age, years, mean (SD) [range] | 68.3 (4.4) [61-75] | 69.7 (4.1) [63-76] | 0.435 |
| Gender, F:M | 9 : 10 | 8 : 9 | 0.985 |
| Education, years, mean (SD) [range] | 12.8 (3.3) [8-19] | 11.1 (3.1) [8-18] | 0.129 |
| Premorbid VIQ, mean (SD) [range] | 110 (7.8) [91-119] | 105 (8.7) [91-123] | 0.083 |
| MMSE, mean (SD) [range] | 28.5 (1.3) [26-30] | 25.5 (2.0) [23-29] | <0.001 |
| Hippocampal volumes, normalized % of TIV*, mean (SD) [range] |  |  |  |
| Right | 0.28 (0.03) [0.20-0.33] | 0.24 (0.03) [0.17-0.30] | <0.001 |
| Left | 0.27 (0.02) [0.23-0.32] | 0.23 (0.03) [0.17-0.28] | <0.001 |
| Total | 0.55 (0.04) [0.47-0.65] | 0.46 (0.06) [0.34-0.58] | <0.001 |
| Biomarkers of Amyloidopathy |  |  |  |
| CSF-Aβ42, mean (SD) [range], cut-off = 700 (n = 7) | - | 569.3 (128.4) [426-735] | - |
| Florbetapir-AV45 (n = 7) | - | All positive | - |

#### 3.1.2. Familiarization and Study phases

Briefly, and importantly, sensitivity (*A_z_*) and omissions (i.e. % Misses) were not different between controls and AD-MCI at immediate recognition of the Familiarization phase (*A_z_*: t=1.478; *p*=0.149; %Misses: t=−0.628; *p*=0.534). Similarly, congruency judgments for face-occupation combinations during the familiarization phase and for face-scene associations during the study phase did not differ between groups (Familiarization, % “Plausible” responses: t=0.836; *p*=0.409; Study, % “Congruent” responses: t=0.176; *p*=0.862). As expected, faster reaction times were observed with repetition within each group during the Study phase in the scanner, despite AD-MCI being overall slower. Importantly, the kind of prior knowledge modulated responses latencies in healthy controls, being faster for PEK over EK, while such interaction remained non-significant in AD-MCI (see **Supplementary Materials 2** for detailed results and statistics).

#### 3.1.3. Test phase

PEK stimuli led to higher item sensitivity and source memory than EK stimuli in healthy controls (Item sensitivity: F(1,18)=9.017; p=0.008; *η^2^*=0.334; source memory: F(1,18)=47.071; p<0.001; *η^2^*=0.723), but not in AD-MCI patients (Item sensitivity: F(1,16)=0.644; p=0.434; *η^2^*=0.039; source memory: F(1,16)=0.446; p=0.514; *η^2^*=0.027), resulting in significant Group x Prior Knowledge interactions (Sensitivity: F(1,34)=5.771; *p*=0.022; *η^2^*=0.141; source memory: F(1,34)=13.05; *p*<0.001; *η^2^*=0.189) (**Figure 2**). Post-hoc analyses using the Holm criterion showed that AD-MCI patients did not differ from controls for EK stimuli for either sensitivity or source memory (Item sensitivity: t=−2.011, p=0.098; Source memory: t=−1.466, p=0.444) but they performed below controls’ level in the PEK condition (Item sensitivity: t=−4.582, p<0.001; Source memory: t=−5.411, p<0.001).

**Figure 2.**
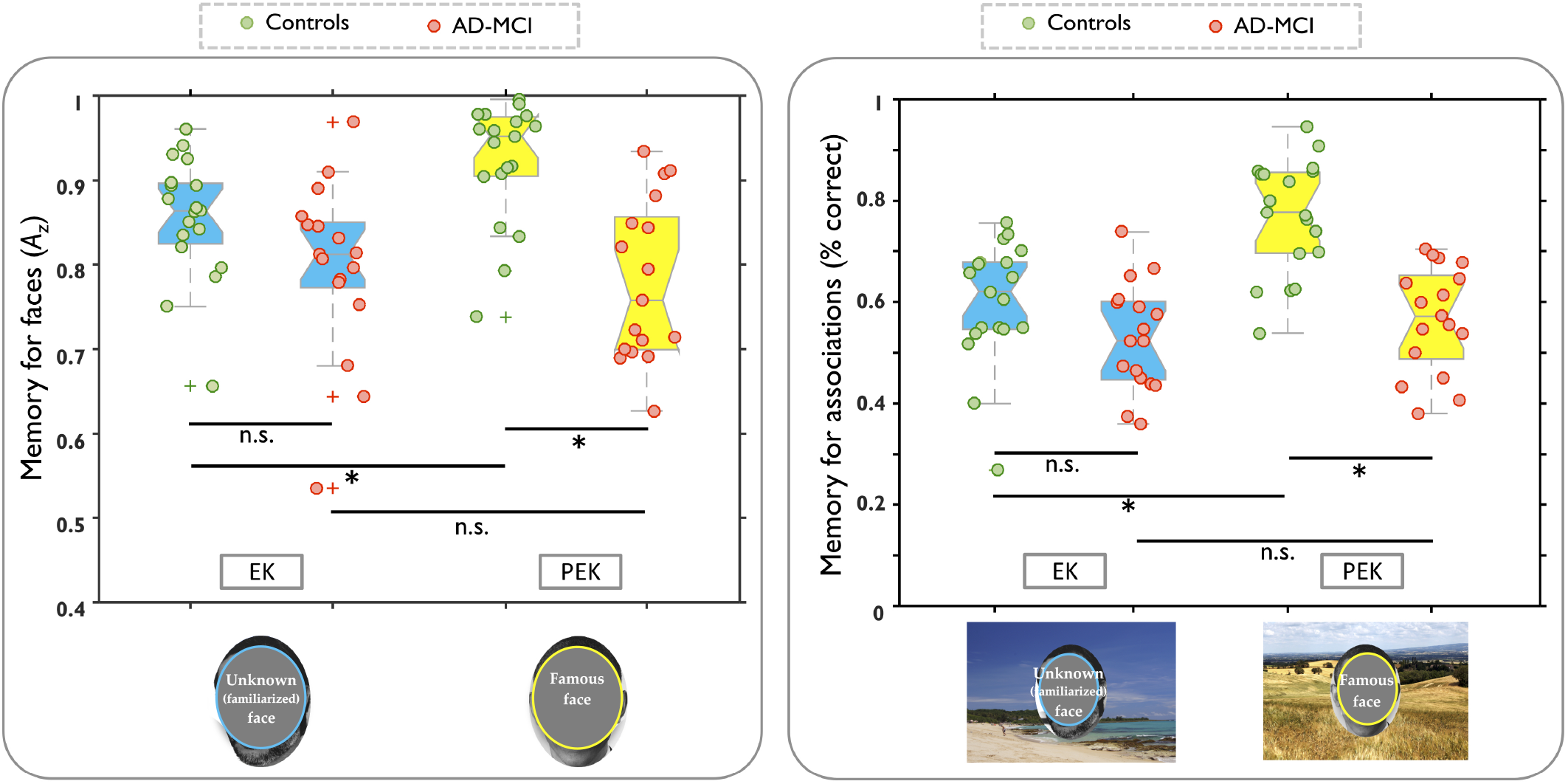
Photographs of faces were obscured due to editor policy. Controls and AD-MCI performance during the Test phase. Left panel: memory for faces; Right panel: memory for face-scene associations. Individual performances (dots) are overlaid on notched boxplots where notches depict the 95% CI around the median. EK: experimentally familiar stimuli (i.e. unknown but familiarized faces); PEK: pre-experimental familiar stimuli (i.e. famous faces). * = p<.05.

### 3.2. Imaging results

#### 3.2.1. Repetition and encoding contrasts

A conjunction analysis between “Repetition” and “Encoding” contrasts yielded activations within occipital and occipito-temporal regions, along the visual ventral pathways (illustration in **Supplementary Material 3**) in both groups. Neural adaptation effects thus allowed accurate identification of the functional networks involved in our visual encoding task.

#### 3.2.2. Prior Knowledge x Repetition interaction

Results for simple contrasts (main effects of Prior Knowledge [i.e. PEK vs EK stimuli encoding] and of Repetition [i.e. Neural adaptation for PEK and EK stimuli repetition]) are provided in **Supplementary Material 4**.

In healthy controls, the interaction contrast [Prior Knowledge x Repetition] revealed activations within three sets of regions (**Figure 3A**): bilateral inferior temporal lobes and occipito-temporal cortices, including MTL structures; bilateral medial and lateral parietal structures; left ventromedial and dorsolateral prefrontal cortices. Multiple ANOVAs revealed four distinct patterns of activations resulting in these interactions. 1) PEK yielded repetition enhancement while on the contrary EK yielded repetition suppression in a number of clusters in bilateral occipito-temporal regions, up to the perirhinal cortex in the right hemisphere but also in the ventral and dorsal prefrontal cortices (**Figure 3A 1**). 2) Parietal clusters showed repetition enhancement for PEK only (**Figure 3A 2**), 3) while bilateral occipital regions showed suppression effects for EK only (**Figure 3A 4**). 4) Last, although PEK was generally associated with repetition enhancement, it was associated with repetition suppression in two small clusters in the posterior and anterior temporal lobes (**Figure 3A 3**). Our results thus support the proposition that in healthy controls, distinct prior knowledge generate opposite neural adaptation in clusters common to PEK and EK, with the addition of partly non-overlapping networks underlying PEK- and EK-based associative encoding.

**Figure 3.**
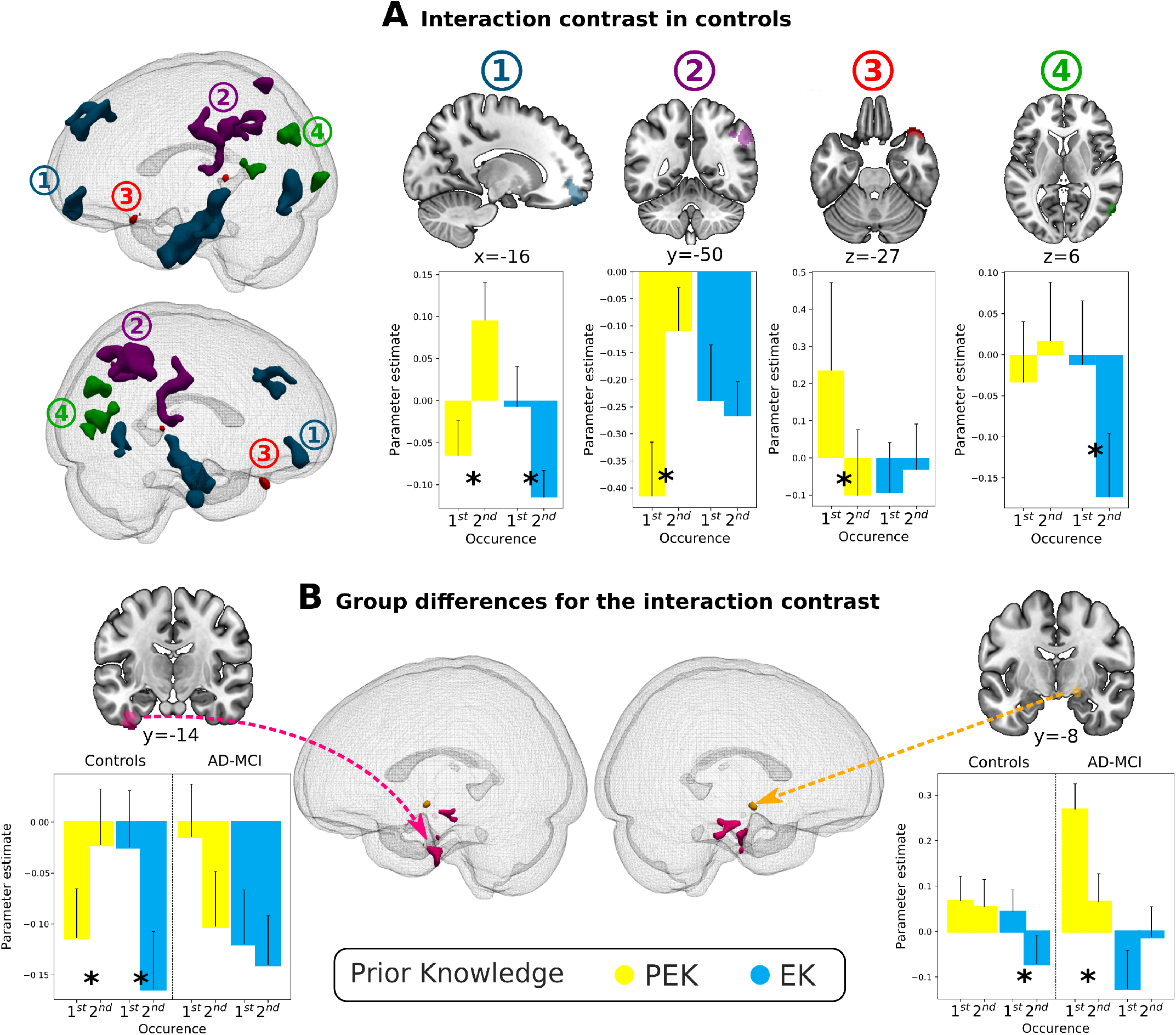
fMRI [Prior Knowledge x Repetition] interaction contrast. (**A**) Distinct patterns of fMRI responses in healthy controls. For each pattern, a plot illustrates the typical neural adaptation effects found within each cluster (coordinates above each plot). For example, the first plot shows that within the left vmPFC, opposite neural adaptation effects were found for PEK vs. EK stimuli, resulting in a significant interaction that was similar within all the clusters of the same blue color displayed in the 3D views on the left. See text for the presentation of the four different patterns. (**B**) Clusters where significant differences were found between healthy controls and AD-MCI patients for the interaction contrast. The clusters in pink show significant neural adaptation effects observed in healthy controls but not in AD-MCI. A cluster in the right hippocampus (in orange) showed neural suppression for EK in healthy controls, but for PEK in AD-MCI. * p<0.05.

In sharp contrast with these results in healthy controls, the same analyses in AD-MCI participants did not yield any significant cluster.

Two samples t-tests yielded differences between the groups for this interaction contrast within two main clusters located in the bilateral inferior temporal lobes (**Figure 3B**). Two additional smaller clusters within the right medial temporal lobe did not reach our clustering threshold, but proved significant after small volume correction for medial temporal lobe structures (as defined with the AAL template, Tzourio-Mazoyer 2002, p<.05; FWE-corrected: right hippocampus (k= 22; main peak x=18 y=−8 z=−12) and right lateral perirhinal cortex (k= 21; main peak x=34 y=−16 z=−36). To investigate these group differences, mixed ANOVAs with Group as between-subjects factor, Prior Knowledge and Repetition as within-subjects factors were conducted on the mean beta weights extracted from the group differences clusters. These analyses revealed the absence of repetition effects in the AD-MCI group, with the exception of the right hippocampus where PEK stimuli yielded repetition suppression. In healthy controls, EK stimuli triggered repetition suppression in the right hippocampus, the right lateral perirhinal cortex and the bilateral fusiform gyri. By contrast, these last two regions (right perirhinal cortex and bilateral fusiform gyrus) showed repetition enhancement effects for PEK stimuli. While these results inform us about the brain activity related to visual associative encoding of face-scene pairs depending on the kind of prior knowledge, they do not speak about the neural substrates of successful associative encoding, which we now report.

#### 3.2.3. Associative Memory effects

Here, we investigate whether the above findings hold for successful associative encoding, i.e. correct retrieval of face-scene associations. We extracted the beta weights associated with the memory events at study within the data-driven ROIs highlighted in the previous section (Source Hits, reflecting accurate associative memory, Source Misses, reflecting accurate item but inaccurate associative memory, and Misses, reflecting face forgetfulness, see Methods section). We then looked for subsequent associative memory effects (i.e. significant differences between Source Hits and Source Misses). First, we did so within clusters derived from healthy controls. Second, we applied the same approach in the clusters derived from the effect of group on the interaction contrast (see previous section). The aim of these analyses was twofold: 1) Does the network involved in associative encoding of PEK vs. EK stimuli also play a role in successful associative memory formation in healthy elderly? 2) Do the regions exhibiting between-groups differences for the [Prior Knowledge x Repetition] interaction contrast also display differential subsequent associative memory effects between groups?

In healthy controls, we found subsequent associative memory effects for both PEK and EK stimuli within the left middle occipital and occipito-temporal areas, as well as within the left vmPFC. A series of regions also showed selective associative memory effects for PEK or EK stimuli respectively (see **Supplementary material 5** for details). Activity in the left DLPFC and in the right medial temporal lobe, including the perirhinal cortex, were higher for PEK Source Hits than PEK Source Misses, but did not discriminate source memory for EK stimuli. Conversely, bilateral precuneus, left fusiform gyrus, left posterior hippocampus, and a right-sided area including the posterior angular gyrus were more activated for EK Source Hits than EK Source Misses. These analyses were not performed in AD-MCI, since no cluster reached significance for the [Prior Knowledge x Repetition] interaction contrast (see previous section).

Finally, subsequent associative memory effects were also found in the regions that responded to the interaction contrast differentially in healthy controls and AD-MCI patients. First, healthy controls and AD-MCI showed similar associative memory effects for EK stimuli in the right hippocampus (albeit only reaching *p*=0.051 in the patient group, **Figure 4A**). However, the right perirhinal cortex and the fusiform gyrus showed higher fMRI response for Source Hits than Source Misses (**Figure 4B**) for PEK stimuli in healthy controls, while this memory effect was not observed in AD-MCI. Moreover, we found that the associative memory contrast estimates (i.e. beta weights difference between Source Hits and Source Misses) in that region correlated positively with behavioral source accuracy measure for PEK stimuli across all participants (r=0.344; *p*=0.02) (see **Figure 4**).

**Figure 4.**
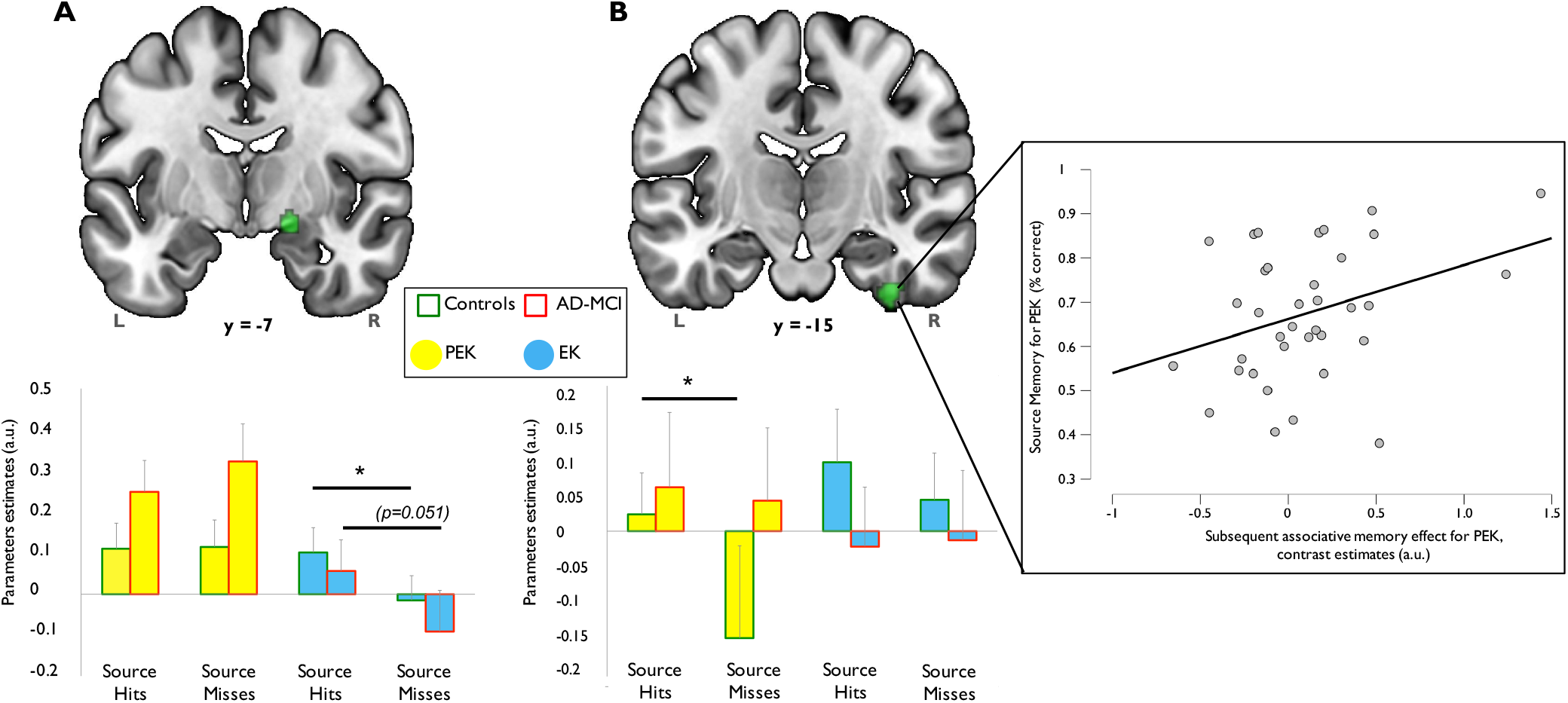
Regions associated with correct face-scene associative memory effects within data-driven ROIs resulting from group differences for the Prior Knowledge x Repetition interaction contrast. (**A**) The right hippocampus showed a similar difference between source hits and source misses for both groups for EK only. (**B**) In contrast, the right perirhinal region showed a difference only for PEK in healthy controls. * p<0.05. On the right, correlation plot showing the association between source memory accuracy for PEK and subsequent associative memory contrast estimates for PEK.

## 4. Discussion

Prior knowledge is recognized as a memory enhancer in healthy subjects, but surprisingly little is known about it in Alzheimer’s disease. In the present study, AD-MCI patients and matched healthy elderly controls learned new face-scene associations. At study, faces were associated with two kinds of prior knowledge. Famous faces (i.e. PEK items) carried prior knowledge resulting from long-lasting multiple exposures, while unknown faces carried prior knowledge due to repeated presentations just prior to study (i.e. EK items). Memory for faces and for face-scenes associations improved for PEK relative to EK stimuli in healthy controls, thus showing the expected effect of prior knowledge. In sharp contrast, AD-MCI patients did not show any benefit of pre-experimental prior knowledge.

In healthy controls, the interaction between repetition and prior knowledge effects on the fMRI response showed the recruitment of large encoding networks partly non-overlapping between EK and PEK. In sharp contrast again, such interaction was absent in AD-MCI patients. Finally, AD-MCI patients displayed impaired activation in a right subhippocampal region where activity predicted successful associative memory formation of PEK stimuli.

### Accounts for the benefits of prior knowledge

The memory advantage for stimuli associated with PEK fits with the levels-of-processing framework (“LOP”, Craik and Lockhart, 1972), in that deeper processing at encoding (i.e. semantic processing of famous faces) results in increased long-term memory formation compared to shallower encoding (i.e. resulting from the absence of, or weaker prior knowledge for an unknown face). In healthy controls, the observation of a larger repetition priming effect on response times for PEK as compared to EK trials at study is consistent with this interpretation.

However, while the LOP explains the PEK advantage for item memory, it does not in itself accounts for the finding of increased accuracy for source memory in healthy controls. Two main accounts have been put forward to explain this “familiarity bonus” on source memory (Poppenk et al., 2010a, 2010b; Poppenk and Norman, 2012). Prior knowledge might reduce the attentional resources required at encoding to build up a new association (Castel and Craik, 2003; Naveh-benjamin and Craik, 1998). Given that we made PEK & EK stimuli familiar to participants at study, this account seems unlikely. Alternatively, an extension of the LOP suggests that famous faces generate more elaborative processing at encoding, enriching events representations (Bein et al., 2015). In turn, enriched representations would become more distinctive and therefore less prone to interference at retrieval. One candidate for this elaborative processing is unitization. It has been suggested that unitization strategies at encoding could alleviate the associative learning deficit in aging (Bastin et al., 2013; Delhaye and Bastin, 2016), which is consistent with our findings of a large associative memory enhancement in elderly controls for PEK vs. EK stimuli.

### The perirhinal cortex is involved in successful pre-experimental knowledge learning

The perirhinal cortex could support some forms of declarative learning, with minimal involvement of the hippocampus, as long as pre-existing representations are available and congruent with the memoranda (e.g. Fernández and Morris, 2018; Kan et al., 2009; Van Kesteren et al., 2012). In healthy controls, we found subsequent associative memory effects for PEK, but not EK stimuli in the right perirhinal area, along with other regions consistently associated with semantic retrieval (e.g. left middle frontal gyrus, Barbeau et al., 2012; Joubert et al., 2010, 2008; Kapur et al., 1994; Martin, 2007; Pineault et al., 2018). Associative memory for PEK, but not EK, was also positively related to the fMRI response in the right perirhinal cortex across both groups.

Such finding could explain the pattern of performance observed in both healthy controls and in AD-MCI patients. The role of the perirhinal cortex in the computational requirements of unitization has already been robustly highlighted in healthy subjects (Diana et al., 2010; Haskins et al., 2008). Semantic retrieval in healthy controls when PEK is involved at study could promote unitization strategies, as supported by perirhinal cortex processing, therefore facilitating the successful formation of new face-scene associations. In contrast, such mechanism could be impaired in AD-MCI patients. Consistent with reports of early person knowledge impairment in the course of AD (e.g. Barbeau et al., 2012; Brambati et al., 2012; Joubert et al., 2010, 2008), activation of pre-experimental semantic knowledge during learning was presumably impaired in our sample of early AD patients, as suggested by our findings of similar repetition priming effects for PEK and EK stimuli in our patient sample during the study phase while they differed in control subjects. Knowledge about unique entities like faces has been shown to be impaired early in AD due to tau pathology in subhippocampal structures (entorhinal and perirhinal cortices) (Braak and Braak, 1995), thus disrupting semantic retrieval (Didic et al., 2011). This interpretation is also consistent with the finding that the advantage of unitization for alleviating the associative memory deficit is absent in early AD patients (D’Angelo et al., 2016), in relation to perirhinal cortex atrophy (Delhaye et al., 2018, 2019).

### Knowledge available at encoding involves different brain networks

More broadly, the interaction between the fMRI responses to neural adaptation and prior knowledge suggests that encoding networks are sensitive to the nature of pre-existing representations associated with the memoranda. Some brain areas displayed opposite repetition effects depending on the kind of prior knowledge involved, extending prior evidence that neural adaptation is not an automatic brain response to stimulus repetition (Henson et al., 2002), rather being associated with either reduced involvement of encoding mechanisms (i.e. repetition suppression), or explicit and/or implicit successful retrieval (i.e. repetition enhancement) (Kim, 2017).

The regions identified in our study as being involved in either types of prior knowledge are consistent with other subsequent memory and / or retrieval studies (ventral and dorsal prefrontal cortices, bilateral occipito-temporal regions, up to the perirhinal cortex; Kim, 2013, 2011; Maillet and Rajah, 2014; Spaniol et al., 2009). We suggest that in the presence of pre-existing semantic knowledge, enhanced activity is observed in regions involved in memory retrieval, schema detection, and visual encoding. However, when recent, episodic-like, pre-existing representations are available, repetition suppression is rather observed in a visual encoding network, including regions involved in the detection of prior occurrence across successive learning trials. For example, we found repetition suppression for EK stimuli along the visual ventral stream, reflecting the reduced engagement of a visual encoding network in the absence of pre-existing semantic knowledge. By contrast, repetition enhancement for PEK stimuli in the same regions may reflect successful retrieval of pre-existing semantic knowledge, along with the involvement of congruency detection processes, as reflected by the involvement of the vmPFC. This region indeed plays a critical role in the detection of congruency between incoming perceptual processing and pre-existing knowledge, or “schemas” (Van Kesteren et al., 2012; Bein et al., 2014). The observation of repetition enhancement for PEK stimuli within the right angular gyrus, with no repetition effect for EK stimuli, also lends support to this interpretation, since this area forms a hub for the representation of prior knowledge (Gilboa and Marlatte, 2017; Wagner et al., 2015). Taken together, our findings therefore underline the dynamic nature of new associative encoding in the elderly, which seems to entail distinct mechanisms along with partly non-overlapping neural networks depending on the kind of prior knowledge involved.

Importantly, this pattern of interactions between prior knowledge and neural adaptation was severely impaired in AD-MCI, and we reported essentially absent or aberrant neural adaptation in this group, in agreement with prior reports (Pihlajamäki et al., 2011, 2008). Therefore, impaired benefit of PEK in AD-MCI patients may be related to both networks and focal (i.e. perirhinal region) dysfunctions that might concur to their poor memory.

### The learning impairment in early AD is likely underestimated

We bring strong evidence that in healthy aging, lifelong accumulated knowledge about a face massively enhances associative learning, beyond what could be expected due to multiple recent exposures (Badham et al., 2015, 2012; Bastin et al., 2013; Umanath and March, 2014). By contrast, we report that AD-MCI patients completely failed to take advantage of such prior knowledge. If replicated, this could open highly promising avenues to better characterize early memory impairments in AD vs. healthy aging, asking whether PEK associative memory is impaired for other types of stimuli and memoranda.

From a practical point-of-view, these results suggest that memory deficits in early AD patients might well be underestimated. When repeatedly presented with unfamiliar stimuli, like in the EK condition, our sample of AD-MCI patients performed fairly well. This situation closely matches the typical multiple-trials learning tasks used in clinical and research settings, where lists of rather unfamiliar items (e.g. isolated words or pictures) are repeatedly presented across learning trials. However, the inability of our patients to take advantage from pre-experimental knowledge would suggest that their ability to do so in everyday routines, where most of the stimuli and events are highly familiar (e.g. neighbors, friends, family met in familiar environments such as shops, streets, etc.), is very likely to be impaired. In other words, while remote prior knowledge increases the likelihood of encoding in healthy aging, it is of no additional value in early AD patients. Still, memory assessment in clinical or research settings usually either do not take into account the role of prior knowledge, or involves familiarization with initially unfamiliar stimuli. One must therefore consider that we might well underestimate the actual memory impairments in early AD patients.

### Conclusion

The present findings bring new evidence for a critical difference in the way early AD patients and healthy elderly form new memories. Pre-experimental prior knowledge proved beneficial for subsequent memory formation in healthy elderly, in relation to the perirhinal region and specific neural networks at encoding. In contrast, impaired benefit of pre-experimental knowledge was observed in early AD, along with abnormal neural adaptation. Such results open new perspectives in our comprehension of the memory difficulties observed in this disease.

## Acknowledgements

We are very grateful to all participants in this study for their time, courage and willingness to advance knowledge. We are grateful to all the MRI operators of the Neurinfo neuroimaging platform for their commitment in public research and for their kindness. We would also like to thank Pierre Gagnepain, Sylvain Charron and Camille Maumet for their valuable advices and availability. MRI data acquisition was performed at the Neurinfo MRI research facility from the University of Rennes I, University Hospital of Rennes, Inria, CNRS and the Rennes Cancer Center. Neurinfo is also supported by the Brittany Council, Rennes Metropole and GIS IBISA.

## Supplementary Materials

### 1. List of the tests used and neuropsychological background

**Table S.1.** Neuropsychological background of the participants

|  | Healthy Controls | AD-MCI | p values |
| --- | --- | --- | --- |
|  | mean (sd) | mean (sd) |  |
| **Global cognition** |  |  |  |
| **MDRS** |  |  |  |
| Global score, max=144 | 141.2 (1.8) | 130.8 (6.5) | <0.001 |
| Attention, max=37 | 36.2 (1.0) | 36.1 (0.9) | 0.696 |
| Initiation, max=37 | 36.7 (0.7) | 31.5 (4.3) | <0.001 |
| Construction, max=6 | 6.0 (0.0) | 6.0 (0.0) | - |
| Concepts, max=39 | 37.7 (1.7) | 37.3 (1.6) | 0.308 |
| Memory, max=25 | 24.6 (0.7) | 20.7 (3.6) | <0.001 |
| **Memory** |  |  |  |
| Logical Memory, WMS-III, Immediate recall | 41.9 (6.8) | 18.1 (7.6) | <0.001 |
| Logical Memory, WMS-III, Delayed recall | 25.8 (5.0) | 3.8 (4.1) | <0.001 |
| DMS-48, pictures, max=48 | 46.2 (2.2) | 41.2 (4.3) | <0.001 |
| DMS-48, words, max=48 | 40.1 (3.9) | 31.9 (4.8) | <0.001 |
| Warrington Memory Test (Faces), max=50 | 41.2 (4.5) | 35.6 (6.5) | 0.005 |
| **Naming** |  |  |  |
| D.O. 80, max=80 | 78.3 (1.5) | 76.2 (3.5) | 0.046 |
| **Limb praxis** |  |  |  |
| Symbolic gestures, max=5 | 4.9 (0.2) | 4.6 (0.6) | 0.056 |
| Tools pantomimes, max=10 | 9.5 (0.8) | 9.1 (1.2) | 0.253 |
| Imitation of abstract gestures, max=8 | 7.6 (0.8) | 6.7 (1.0) | 0.003 |
| **Executive functions** |  |  |  |
| Trail Making Test Part A (seconds) | 33.5 (9.0) | 42.5 (10.0) | 0.007 |
| Trail Making Test Part B (seconds) | 70.2 (21.6) | 133.0 (55.9) | <0.001 |
| Trail Making Test Part B (errors) | 0.5 (1.0) | 0.9 (0.9) | 0.043 |
| Stroop test |  |  |  |
| Naming (seconds) | 61.1 (8.1) | 79.6 (16.4) | <0.001 |
| Naming (errors) | 0.6 (1.2) | 2.3 (2.7) | 0.009 |
| Reading (seconds) | 43.3 (5.0) | 48.4 (7.7) | 0.045 |
| Reading (errors) | 0.3 (0.7) | 0.06 (0.3) | 0.339 |
| Interference (seconds) | 106.7 (14.0) | 165.4 (65.2) | <0.001 |
| Interference (errors) | 1.5 (3.3) | 4.8 (4.2) | 0.003 |
| Flexibility (seconds) | 123.1 (20.3) | 211.8 (80.8) | <0.001 |
| Flexibility (errors) | 5.4 (5.4) | 11.2 (6.8) | 0.002 |
| Verbal Fluency |  |  |  |
| Letter "R", 90 seconds, nb. Correct | 20.4 (5.4) | 15.6 (5.5) | 0.012 |
| Category "Fruits", 90 seconds, nb. Correct | 17.4 (3.7) | 12.7 (4.0) | <0.001 |
| **Anxiety & Depression** |  |  |  |
| Spielberger STAY-B, raw score | 35.5 (6.9) | 38.9 (8.7) | 0.214 |
| Beck Depression Inventory-II, raw score | 4.8 (5.7) | 7.4 (6.6) | 0.296 |
| **Subjective forgetting** |  |  |  |
| Memory Self-Evaluation Questionnaire, mean score | 2.2 (0.5) | 2.7 (0.5) | 0.003 |

### 2. Behavioral results from the familiarization and study phases

**Table S.2.**
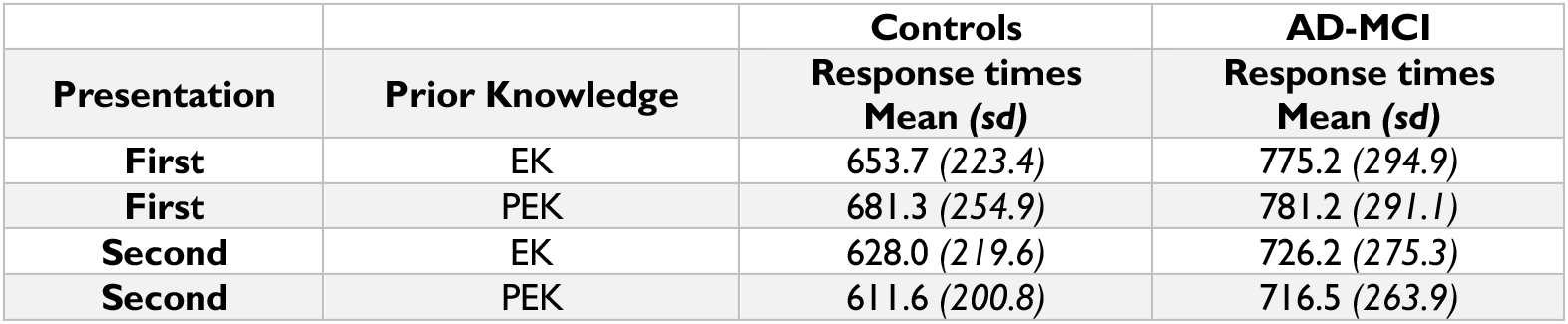
Response times during the study phase

**Figure S.1.**
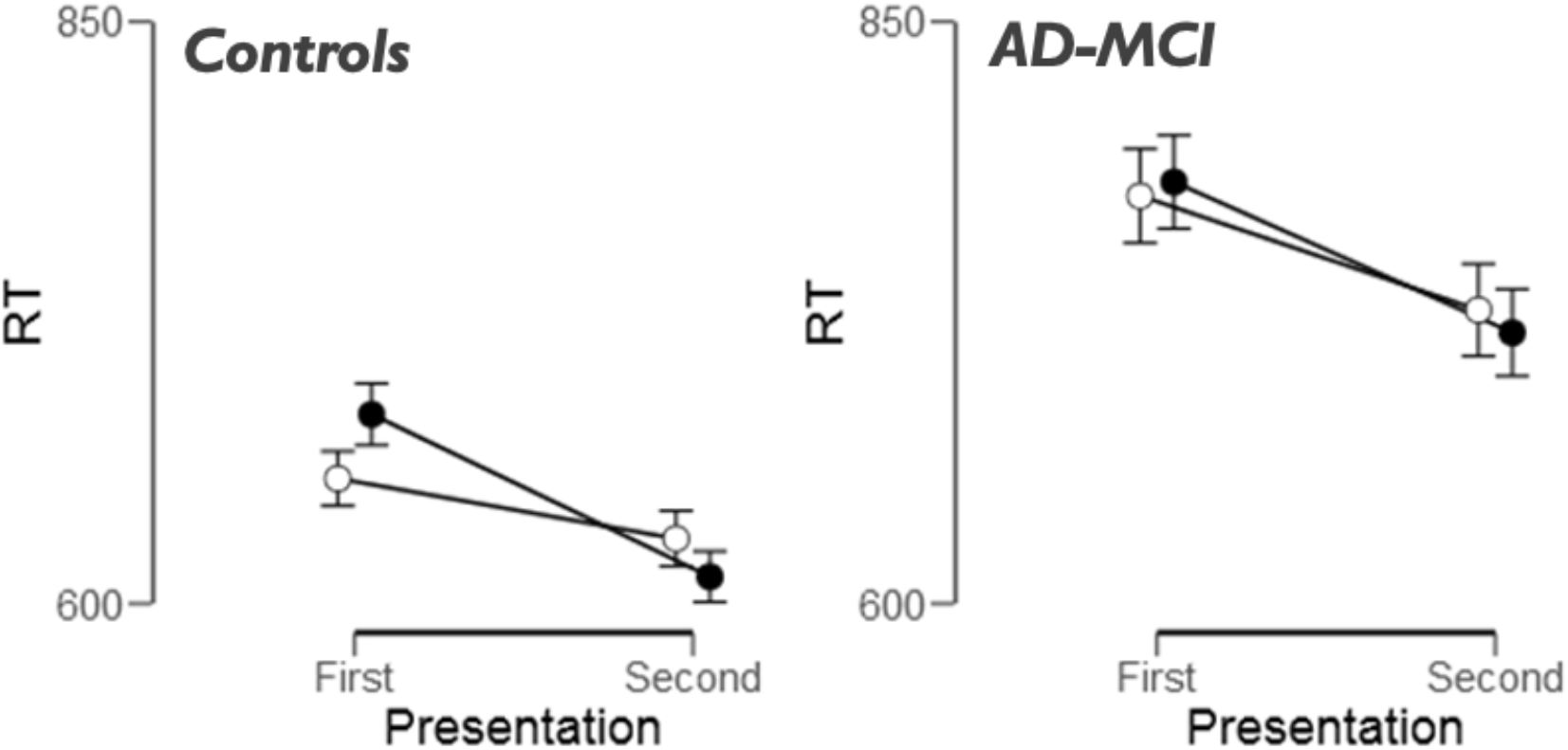
Repetition priming effects on Response Times in Controls (left) and AD-MCI (right) participants (study phase). Black circles = PEK; White circles = EK. Error bars show %95 IC around the mean. AD-MCI displayed slower RTs overall, but presented a significant priming effect (i.e. RT fastening for both PEK & EK stimuli at second presentation). However, the amplitude of priming in Controls was higher for PEK than for EK stimuli (significant PK x Repetition interaction, F=13.388; p<0.001; η^2^=0.014), while it did not differ with the kind of PK in AD-MCI (non-significant PK x Repetition interaction, F=0.667; p=0.414; η^2^=0.001).

### 3. Conjunction analysis: Repetition Suppression and Encoding contrasts

**Figure S.2.**
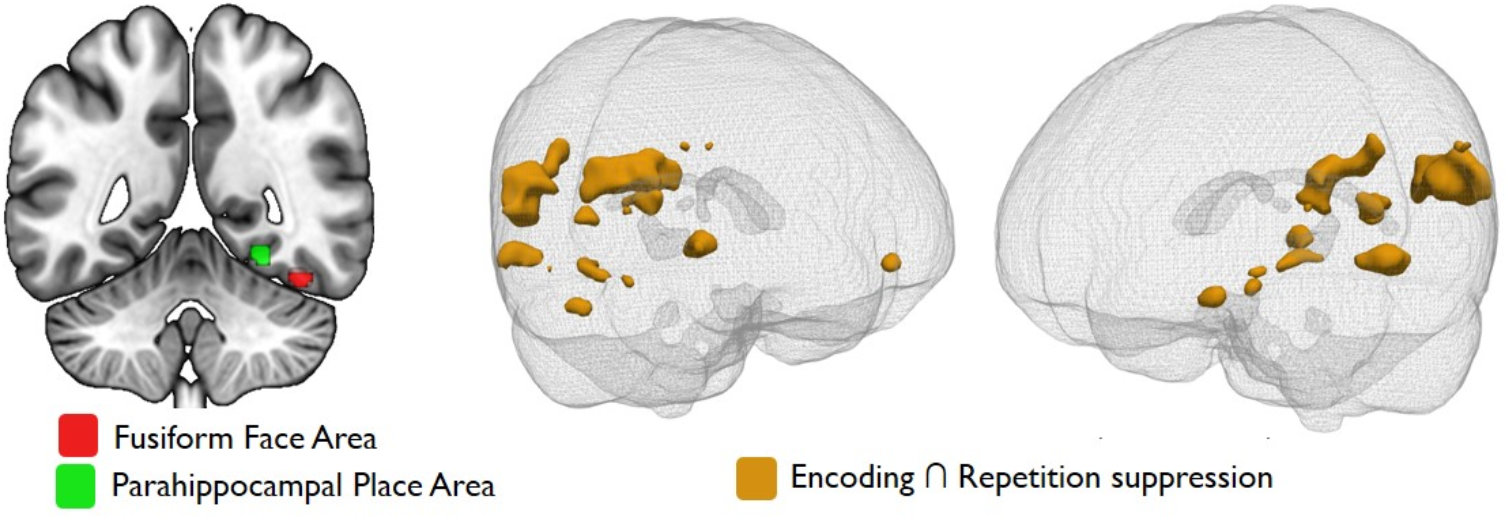
Significant overlaps between {Encoding} and {Repetition suppression} contrasts maps in Controls, p<0.05 FWE corrected. The right-sided 3D view illustrates the significant clusters from the conjunction analysis; the left-sided coronal view focuses on two subregions of the conjunction map, located in the vicinity of FFA and PPA. Note that in AD-MCI patients, the same analysis yielded activation in the Left Parahippocampal Place Area only, albeit at an uncorrected p=0.005 threshold (k=19 voxels, peak MNI coordinates: x=−26; y=−48; z=−12).

### 4. List of supra-threshold clusters for the main effects of Prior Knowledge and Repetition

**Table S.4.**
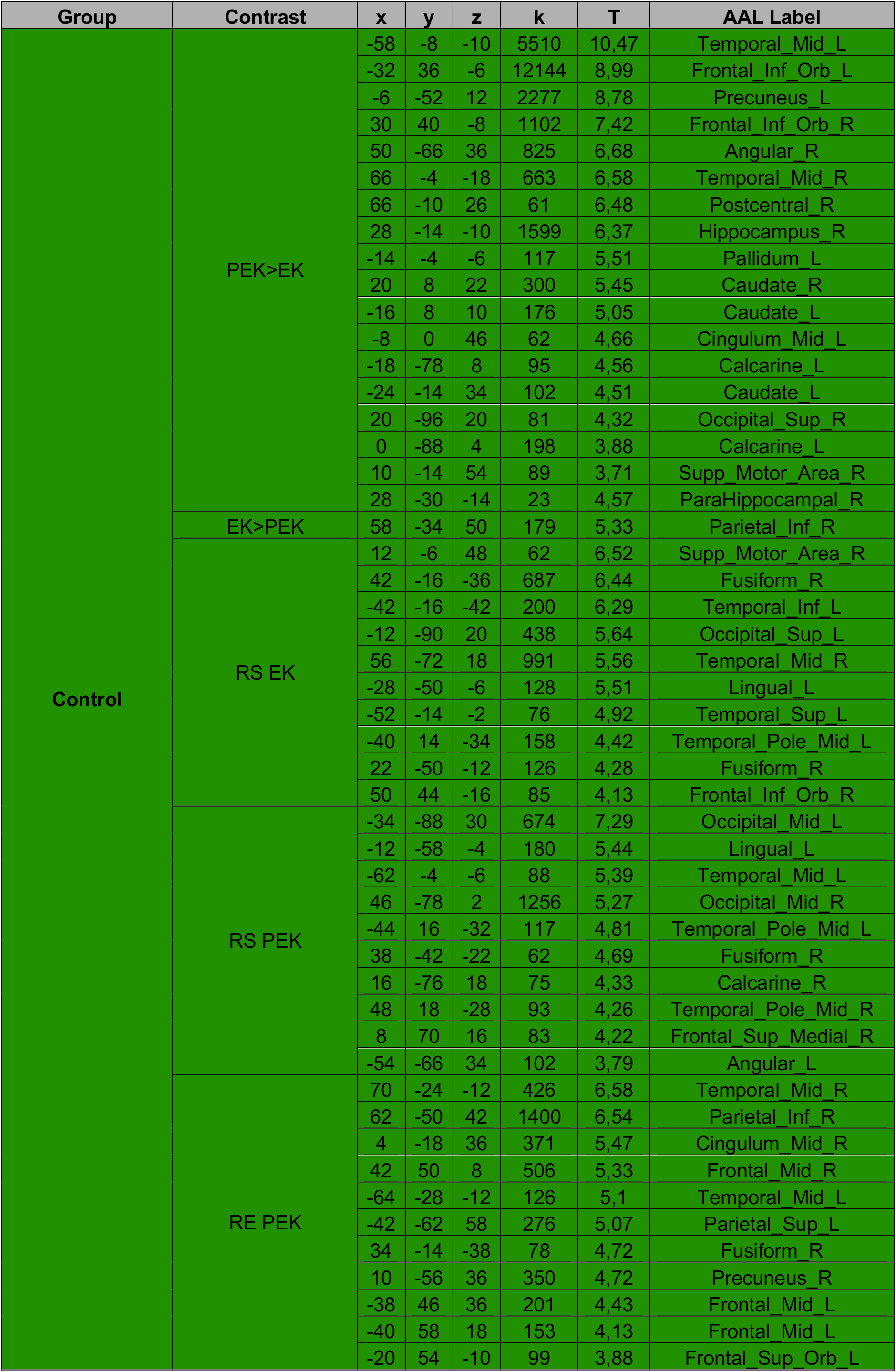
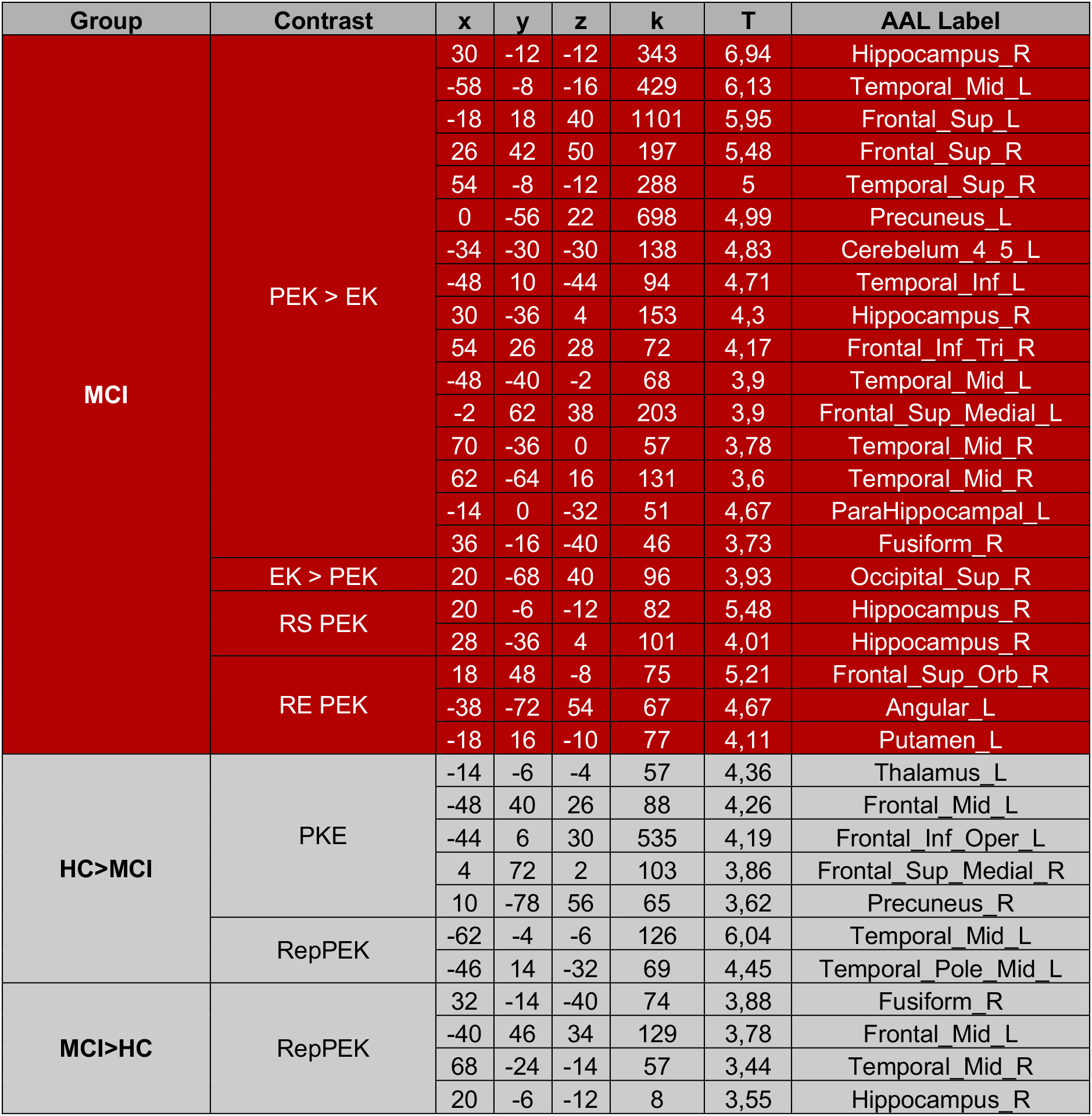
List of suprathreshold clusters for the main effects of Prior Knowledge (“PKE”= Prior Knowledge Effect, PEK vs. EK) and Repetition (Rep; RE=Repetition Enhancement, i.e. more fMRI response to the second over the first presentation; RS=Repetition Suppression, i.e. more fMRI response to the first over the second presentation), for Controls (Green background), AD-MCI (Red Background) and group differences for these contrasts (Grey background).

### 5. Illustration of the subsequent associative memory effects in Controls

**Figure S.3.**
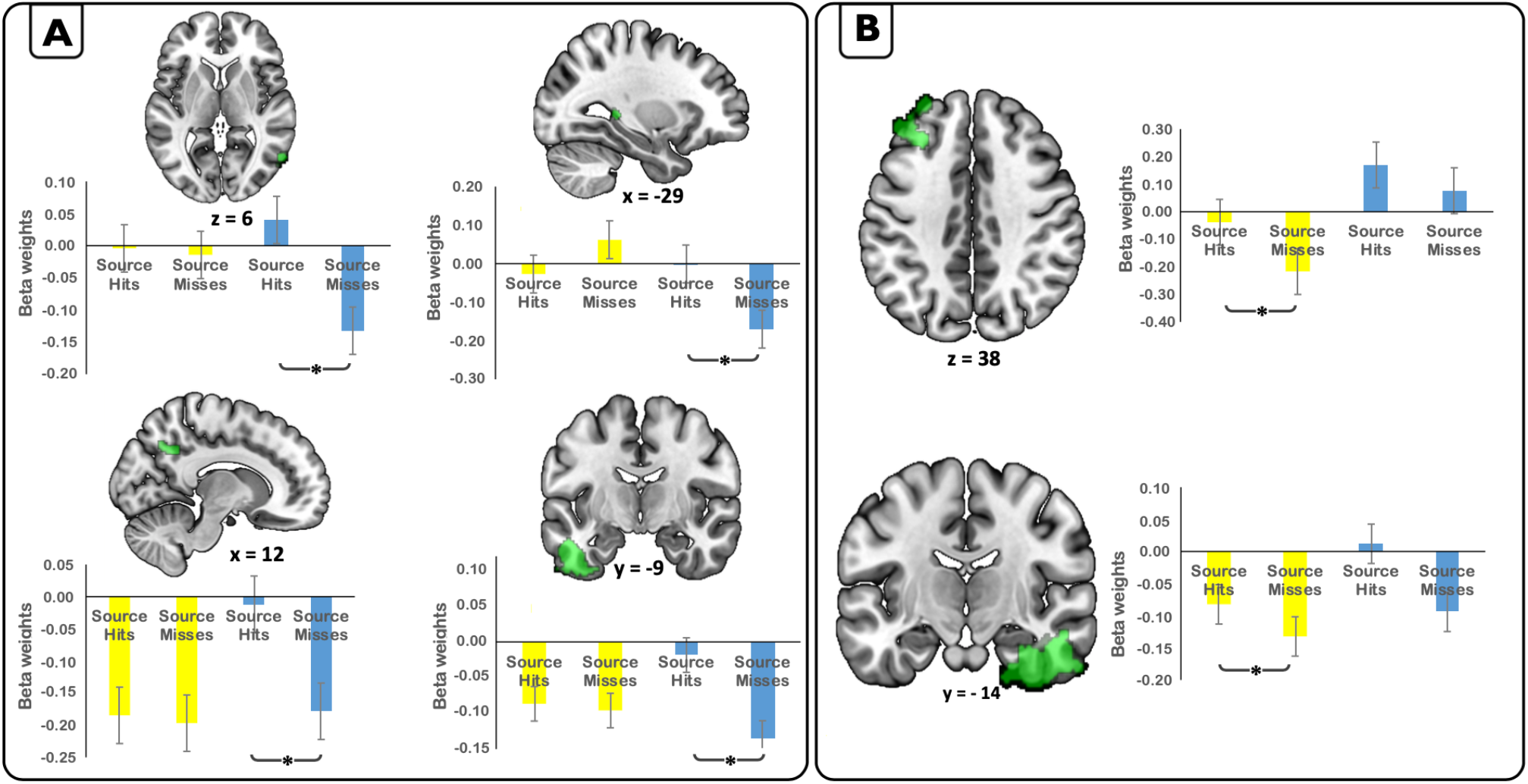
Subsequent associative memory effects (i.e. higher activity for Source Hits than Source Misses) within data-driven ROIs resulting from the Prior Knowledge x Repetition interaction contrast in Controls. (**\****p* values < 0.05). (**A**) Subsequent associative memory effects significant for EK stimuli only; (**B**) Subsequent associative memory effects significant for PEK stimuli only. Regions showing associative memory effects for both PEK and EK stimuli are not displayed (see main text).

